# The Social and Nutritional Factors Controlling the Growth of Honey Bee (*Apis mellifera*) Queens

**DOI:** 10.1101/2024.09.05.611439

**Authors:** Omer Kama, Hagai Y. Shpigler

**Affiliations:** Department of Entomology, Agricultural Research Organization, The Volcani Institute, Rishon LeZion, Israel; The Robert H. Smith Faculty of Agriculture, Food and Environment, The Hebrew University in Jerusalem, Rehovot, Israel

**Keywords:** Honey bees, Honey bee queen, Grafting, Lab assay

## Abstract

The honey bee queen is essential for colony function, laying thousands of eggs daily and determining the colony’s genetic composition. Beekeepers cultivate and trade queens to enhance colony health and productivity. Despite its significance, artificial queen rearing in foster queenless colonies has remained largely unchanged for over a century, offering limited control over the environmental conditions influencing larval development. In this study, we developed a laboratory-based method for queen bee rearing, establishing a protocol for rearing queens in cages in the lab under controlled environmental conditions. We first investigated the minimal number of worker bees required to rear a single queen and found that groups of 200 workers raise queens with comparable success and weight to those reared in colonies. As a proof of concept, we examined the impact of larval age on rearing success in our new system. We found that the age of the larvae affects the success rate of queen development and that younger larvae developed into heavier queens than older larvae, as published in the past using the traditional queen-rearing method. Additionally, we assessed the influence of pollen nutrition on queen-rearing success, finding that a high pollen concentration is crucial for queen development. These findings and the new method provide a foundation for studying queen bee development in a controlled environment and offer potential applications for improving queen-rearing practices.

## Introduction

The honey bee (*Apis mellifera*) queen is the most important individual within the colony, as the reproductive success of the colony depends on her viability (1). Beekeepers typically replace their queens at least once every other year to maintain colony vigor (2). To meet this demand, new queens are raised annually. Artificial queen rearing was developed in the late 19th century in the U.S. by Gilbert Doolittle (3, 4), shortly after the establishment of the modern Langstroth method of beekeeping. This process is based on the principle that every diploid egg has the potential to develop into either a queen or a worker, depending on the social and physical environment of the developing larva (5). A new queen can be artificially reared by transferring young larvae from worker cells into large, vertically oriented queen cells in queenless colonies a process known as grafting (3). Queen breeders, utilizing specialized equipment and knowledge of honey bee biology. The queen-rearing industry includes thousands of breeders who grow millions of queens annually (6).

The honey bee queen has unique morphology, physiology, and behavior that support her tasks performance in the hive with maximal efficiency (1). Throughout her life cycle, the queen’s behavior and physiology adapt to meet the demands of each stage. In the early stages of her life, after emerging from the pupa, the queen displays high aggressive behavior towards rival virgin queens, coinciding with the development of her venom sac and stinger. During her second week, the virgin queen embarks on nuptial flights, during which her flight muscles and navigation abilities fully develop (7, 8). Once the queen returns from her mating flight, she remains in the hive, dedicating most of her time to egg-laying, venturing outside only during swarming events (9, 10). At this stage, her reproductive system and ovaries are fully developed, while her flight abilities and venom sac diminish. The success of a high-quality queen throughout all life stages depends significantly on her genetic background and the environment in which she was raised (11, 12).

Honey bee colonies will rear new queens under three primary circumstances: in spring before swarming, as a replacement for a malfunctioning queen, or at emergence when the queen of the colony unexpectedly dies (1). The queen’s development time is shorter than worker bees (10). Early emergence from the pupa is advantageous as it minimizes the period the colony is queenless, which is crucial for maintaining the colony’s reproductive potential (13). Moreover, Competition among young queens is intense; the first queen to emerge typically attacks and kills rival queen pupae by stinging them before they can emerge (1). Queen breeders often raise multiple queens simultaneously within a single colony. To protect the pupae from the aggression of other queens, breeders separate the pupae around day ten of their development. A foster colony used for queen rearing typically contains tens of thousands of workers, frames with pupae, no young larvae, and no queen. The colony is well-nourished with honey and pollen to support the growth of queen larvae. Breeders usually introduce 40-60 larvae per cycle into the colony (1, 3). The queenless colony will then rear these larvae into queens as an emergency response. To ensure high-quality queens, breeders carefully construct the best possible foster colonies and select larvae from superior colonies. This method is highly efficient and meets market demands. However, breeders have no control over the nutrition that the colony gathers, and the foster colonies are exposed to environmental factors such as weather changes, predators, parasites, pesticides, and diseases including viruses (14). As a result, the rearing process sometimes fail.

The quality of a honey bee queen is influenced by both internal and external factors. Internally, it depends on the genetic background of the larvae and the genetic source of the workers that rear the queen (12). Externally, environmental factors such as the age of the larvae, the type of nutrition provided, and the strength of the foster colony play critical roles (3, 12). Former studies shows that younger larvae, develop into larger and more fertile queens compared to older larvae (15). However, the control that a queen breeder has over the growth process is limited and generally ends after the larvae are introduced to the foster colony. In contrast, bumble bee breeding is conducted indoors, where conditions are tightly controlled to maximize success (16-18). This indoor breeding approach frees bumble bee breeders from the constraints of seasonality, enabling the year-round production of pollinators. However, this approach is not feasible for honey bees, since they live in large perennial colonies that cannot thrive indoors. This limitation reduces the ability of breeders to control environmental conditions, nutrition, and exposure to pathogens during the honey bee queen’s development. Controlling the rearing conditions of honey bee queens can improve their quality and accelerate selective processes

In the current study, we introduce a new method for queen rearing under controlled environmental conditions. This method involves rearing queens using groups of bees housed in cages within a controlled environmental chamber, with a single larva introduced to each group for queen rearing. The method allow the conduction of large experiments with repeatable and reliable replications. Our first experiment aimed to determine the minimum number of bees required to successfully rear a queen. We found that two hundred bees can reliably accomplish this task, but one hundred are not enough. We applied this method to investigate several research questions. As a proof of concept, we tested the effect of larval age on the development of queens within our system. As expected, we found that younger larvae developed into heavier queens as reported in the literature (19-21). Lastly, we explored the impact of nutrition on larval acceptance rates and the quality of the developing queens. By providing the bees with a pollen cake containing an elevated concentration of pollen, we demonstrated the significant effect of nutrition on queen growth. This new method and the findings presented here open the queen-rearing process to detailed investigation under controlled conditions, enabling further study of various factors affecting queen growth.

## Materials and Methods

### Bees

Bees from the apiary of the Volcani Center in Israel were used for the study. The experiments were conducted in the summer of 2023 and spring of 2024. Frames with emerging pupae were collected from the colonies and placed in a six-frame hive box in an acclimated room (34ºC ± 1, RH = 60% ± 5). The day-old emerging bees were then collected into a plastic box and transferred to cages. The number of bees in each cage was estimated by weight using a scale with an accuracy of 0.01 g.

### Cages

We used queen monitoring cages for this study (22). The cage size is 10 × 15 × 5 cm, with the back of a plastic frame. Each cage has four round holes designed to fit 5 ml tubes (diameter – 9 mm), two on the sides and two at the top. These holes are used for feeding and for the introduction of larval queen cups. The cages were supplied with pollen paste (90% pollen / 10% sugar water, 60%), a 5 ml tube of honey, and a 5 ml tube of water. The food was refreshed every other day. The cages were kept in an acclimated room with conditions resembling a normal hive (Temp: 34° ± 1°, RH: 60% ± 5%) for the entire experiment.

### Queen rearing

Frames with young larvae were collected from the hives. Day-old larvae were collected from the cells using a grafting tool and gently transferred into queen-rearing cups (JZBZ queen bee cells) with a small droplet of royal jelly. The cups were introduced to the groups of bees in the cages through a hole at the top of the cage, one cell per cage. For control, cups from the same grafting session were introduced to a foster colony containing six frames of bees and no brood. The cells were monitored to estimate the success rate of larva acceptance after 72 hours, queen cell cup capping, and the rate of fully developed mature queens.

### Queen weight

Since the queen’s weight changes during her adult life, we weighed the queen at the pupa stage when the environment does not affect her weight. The queen pupae were gently removed from the queen cells on day 10 after grafting, approximately one to two days before emergence. The pupae were weighed on an analytical scale to the nearest milligram to estimate the queen’s weight. The pupae were then gently inserted back into the queen cups and sealed with wax. In experiment 2 (see below) the cups were left in the acclimated room for emergence, with each pupa in a separate cage. This manipulation did not affect the hatching success of the pupae. The emerging queens were collected and frozen at -20°C for further analysis.

## Experiment 1: The Effect of the Number of Workers on Queen Development

The cages were populated with different numbers of workers. The number of workers in each cage was estimated by weighing the day-old bees added to the cage, with an estimation of ten bees per gram. The cages were populated with 5 g, 10 g, 20 g, or 30 g of bees representing 50, 100, 200 or 300 bees respectively. One day old larva, placed in a queen cup, was introduced to each cage and tracked for development. Three stages of development were monitored: larval acceptance after 3 days, the number of capped cells after one week, and queen emergence after 14 days. The acceptance and development of the larvae, as well as the weight of the queens, were measured as detailed above. The experiment was conducted using two different colonies.

## Experiment 2: The Influence of Larval Age on Queen Development under Lab Conditions

Larvae of three different ages were used for queen rearing in cages. The cages were populated with 20 g of bees and fed *ad libitum* with honey and 90% pollen cake. To collect larvae of known age for grafting, a honey bee queen was caged in a queen excluder cage on a frame in a regular colony for 12 hours. The frame was checked for eggs at the end of the caging period, and the queen was released. The frame was kept in the cage for three days to ensure that the queen did not lay any more eggs on it. After three days, the frame was removed, and 12 larvae, aged 1-12 hours old, were used for queen rearing under controlled conditions. The frame was then returned to the queen excluder cage in the colony. A day later, on day four, the same frame was used for a second grafting of 24-36 hour-old larvae and returned to the colony. A final round of grafting was performed from the same frame for 48-60 hour-old larvae the next day. The acceptance and development of the larvae, as well as the weight of the queens, were measured as detailed above. The experiment was done with two different colonies.

## Experiment 3: The Influence of Pollen Nutrition on Queen Development

The richness of the pollen cake was varied between the cages to test the effect of pollen nutrition on queen-rearing success. The bees were fed pollen paste at four different levels of richness: Sugar water (60%) with no pollen (P0%), 30% pollen and 70% sugar water (P30%), 60% pollen and 40% sugar water (P60%), 90% pollen mixed with 10% sugar water (P90%). The cages were supplied with *ad libitum* honey as a carbohydrate source. One day-old larva was grafted and introduced, one to each cage. The acceptance and development of the larvae, as well as the weight of the developing queens, were measured as detailed above. The experiment was done with two different colonies.

## Data Analysis

The chi-square test of independence was used to compare the success rate in larval acceptance and queen development. A post hoc analysis was performed using a chisquare test of independence for each pair with FDR correction for multiple comparisons. The weight the queens was compared using the ANOVA test. The data analysis for this study was conducted using the Real Statistics Resource Pack software (Release 8.9.1). Copyright (2013 – 2023) Charles Zaiontz. www.real-statistics.com.

## Results

### Experiment 1: The Effect of the Number of Workers on Queen Development

We compared the queen-rearing success rate in groups of bees with different numbers of workers. The acceptance of the larvae was affected by the number of workers in the group. Fifty bees (50W) accepted only 25% of the larvae and did not grow any of the larvae to a mature queen (n = 20). Groups of 100 bees (100W) accepted 55% of the larvae on day three and grew 20% of them to maturity (n = 20). Two hundred bees (200W) accepted 69% of the larvae and grew 48% to queens (n = 19). Three hundred bees (300W) accepted 84% of the larvae and grew 63% to maturity. (Fig.1A, χ^2^ test for independence, χ^2^_(6)_ = 24.2, p < 0.001). The success rate of queen-rearing in a control colony was 66% (n = 20), which was not different from the success rate of the 200W or 300W groups. In a post hoc analysis, we found the success rate of 50W is significantly lower than that of 200W and the 300W groups, and lower but not significant than the 100W (χ_2_ test with FDR correction, p < 0.05). The 100 group-reared queens were not different from the two and the three hundred groups (χ_2_ test with FDR correction, p > 0.05).

The weight of the queens was also affected by the group size. The queens from the 100W group weighted 194 mg ± 1.0 (n = 2), queens of the 200W group weight 228 mg ± 3.8 (n = 8), and queens from the 300W group weight of 218 mg ± 2.6 (n = 10). The control colony raised queens with an average weight of 236 mg ± 2.6 (n = 13). The differences between the groups were significant (Fig 1B, One-way ANOVA, F_(3)_ = 14.7, p < 0.001). In post hoc analysis we found a significant difference between the 100W group and all other groups (Tukey’s post hoc test, p < 0.05). The colony’s queens were heavier than the 300W queens (p < 0.05) but not significantly different from the 200W queens. The 200W queens were similar in weight to the 300W queens.

**Fig 1:**
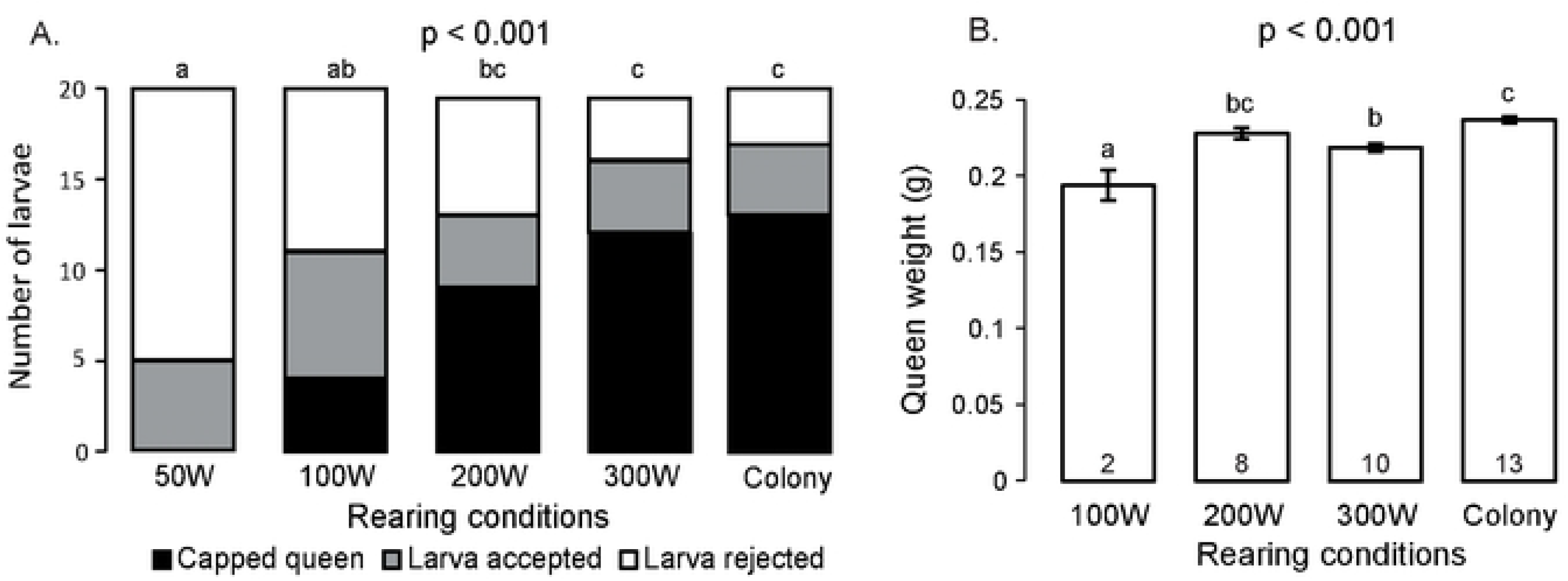
The effect of group size on queen development. A. The success rate of queen growth by groups of bees of different sizes. The column represents the number of larvae rejected by the bees (White) accepted as larvae but were not caped and pupated (Grey) capped queen pupa (Black). The *p-value* is a result of a χ^2^ test of independence, Columns with different letters are significantly different in post hoc test (χ^2^ test with FDR correction p < 0.05). **B**. The weight of queen pupae of grown by groups of bees of different sizes or by a full colony. The column represents the average ± SE; the *p-value* is a result of the One-way ANOVA test. Columns with different letters are significantly different in the Tukey pos-hoc test (p < 0.05).

### Experiment 2: The Influence of the Larvae Age on Queen Development in Cages

We tracked the acceptance and rearing success of larvae at three different ages. We used groups of 200 bees for this experiment. The accepted rate for young larvae at the age of 0-12 hours was 90%, and 75% developed to mature queens (n = 20); larvae at the age of 24-36 hours were accepted at a rate of 96% and 68% were fully developed (n = 25). Only 24% of the 48-60-hour-old larvae were accepted and 16% developed to queens (n = 25). The age of the larvae has a significant effect on the success rate of the development to queens (Fig 2A, Chi-square test for independence, χ_2(4)_ = 37.3, p < 0.001). In a post hoc analysis, we found the success rate of the 0-12h group and the 24-36h are higher than the 48-60h group (χ^2^ test with FDR correction, p < 0.05). The 0-12h group is not different from the 24-36h in the success rate (χ^2^ test with FDR correction, p < 0.05).

**Fig 2:**
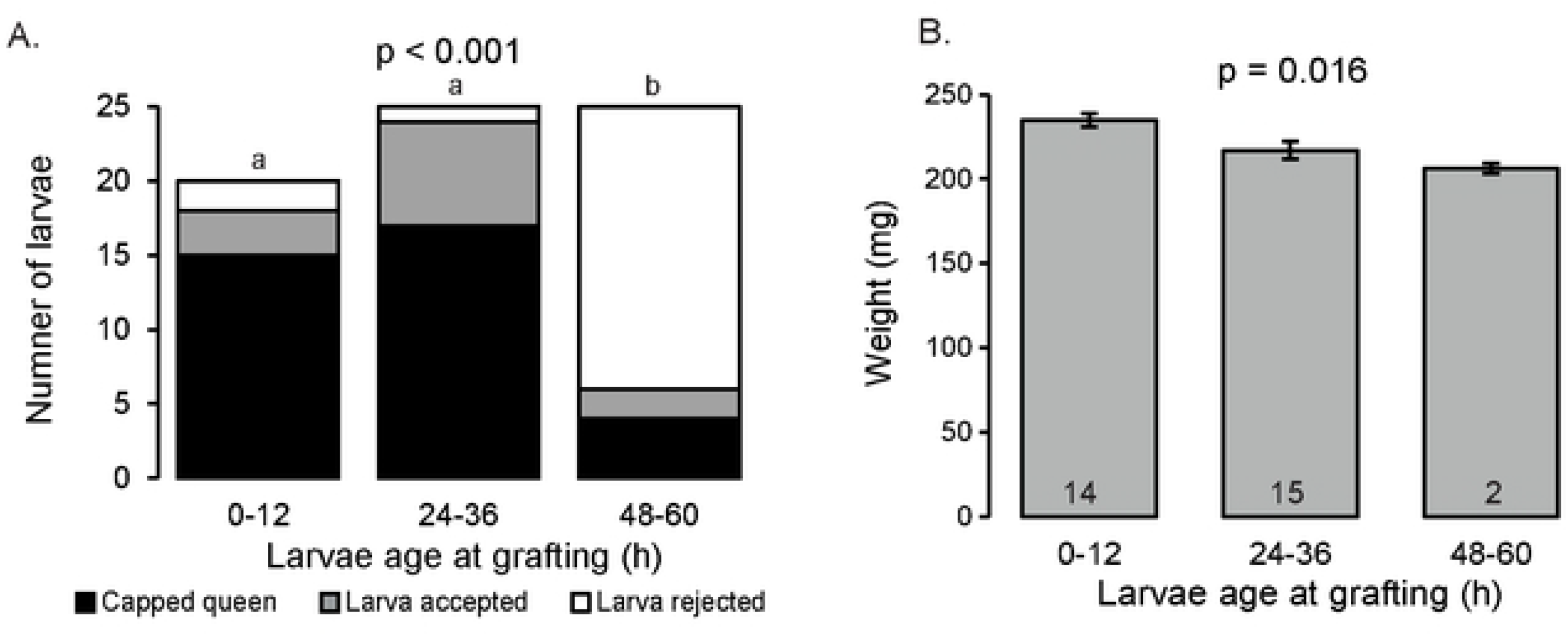
The effect of larval age on queen development under controlled environment. The success rate (A) and the weight (B) of queen reared by 200 bees from larvae from the same source at three different ages: 0-12 hours, 24-36 hours, 48-60 hours. Detailed as at Fig 1.

The weight of the queens was affected by the age of the larva. Queens developed from 0-12h larvae weighed on average 234 mg ± 5 (n = 14) while 24-36h old weighed only 217 mg ± 6 (n = 15), the 48-60 h old larvae developed queens weighed only 206 mg ± 1 (n = 2). The difference between the groups is significant (Fig 2B, One-way ANOVA, F_(2)_ = 4.8, *p* = 0.016). In the Tukey post hoc test, the 0-12h larvae were heavier than the 24-36h larvae (*p =* 0.03). There is no significant difference between the queens developed from the 24-36h and the 36-48h old larvae. Nonetheless, the sample size of the 36-48h group is low due to the low success rate of queen development in this group.

### Experiment 3: The influence of pollen nutrition on queen development

To test the effect of nutrition, we measured the success of queen-rearing by bees fed with pollen paste at different enrichment levels. We used groups of 200 bees for this experiment. We tested four levels of pollen paste enrichment (including pollen and sugar water): No pollen (P0%), P30%, P60% and P90% pollen paste. Removing the pollen from the cages resulted in zero acceptance of the larvae by the workers, and on day three, all the queen cups were empty and clean. Using 30% pollen paste, the bees accepted only a single larva out of 25 (4% success). At medium pollen paste of P60%, the bees accepted four out of 25 larvae (16%), and only two were reared to an adult queen. Using the heavy pollen paste of 90%, eighteen out 25 (72%) of the larvae were accepted by the workers on day three, and 16 were raised to mature queens. The difference between the groups is significant (Fig 3A, Chi-square test for independence, χ^2^_(6)_ = 50.8, *p* < 0.001). In a post hoc analysis, we found the success rate of the P90% group higher than all other groups (χ^2^ test with FDR correction, p < 0.05). The P0%, P30% and P60% are not different from each other in the success rate of queen rearing (χ^2^ test with FDR correction, p > 0.05).

**Fig 3:**
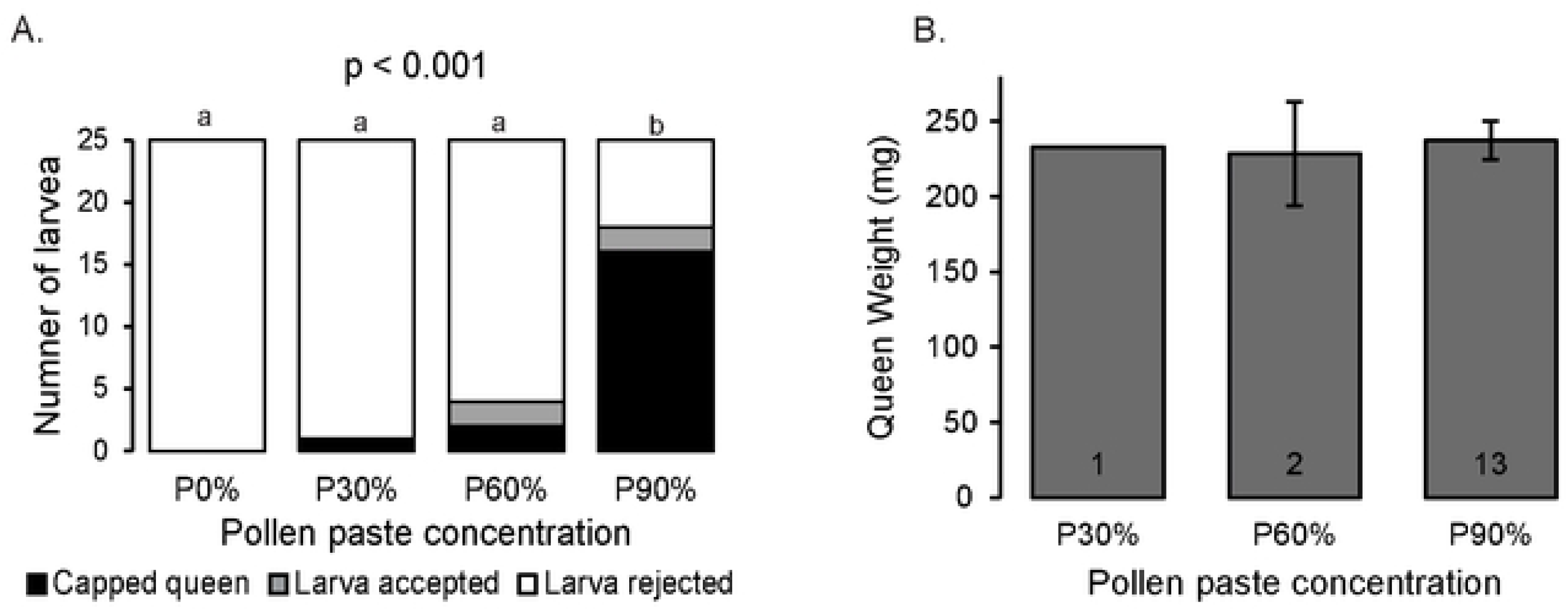
The effect of pollen nutrition on queen development. The success rate (A) and the weight (B) of queen growth by groups of 200 bees fed with different pollen paste concentration. Detailed as at Fig 1.

The average weight of the queen in the P90% group was 237mg ± 4 (n = 13). The average weight of the queens in the P60% group was 228mg ± 2 (n = 2), and the single queen developed at the P30% group weight was 233mg (Fig 3B). Due to the low rearing rate of the queens in the low pollen groups, the differences between the groups were not compared.

## Discussion

The process of queen development depends on various biotic and abiotic factors (23-25). Queen rearing is a social effort, and the workers’ decision to rear a queen is made collectively. In this study, we developed a new protocol to investigate the environmental and social conditions required for successful queen development. Our findings demonstrate that the workers’ decision to rear a queen is influenced by several social and physiological factors, including the number of bees in the group, their nutritional state, and the age of the larvae.

We found that a minimum number of worker bees is necessary to successfully rear a queen from a day-old larva. Fifty workers accepted only a few larvae and nurtured them until day three, but none successfully reared a queen. One hundred bees accepted about half of the larvae, but only a few were raised to maturity. Groups of 200 or 300 workers succeeded in rearing queens at rates not significantly different, compared to a full colony in the current study and consistent with data from other studies (3, 26-28). We assessed queen quality using wet weight of the queens, a common metric for predicting queens’ reproductive potential and hive performance (11, 29-33). The weight of queens reared in the lab by 200 bees was comparable to that of queens reared in traditional colony-based systems. This finding suggests that the indoor queen rearing method developed in this study is comparable in quality to colony-based rearing and that 200 bees is the minimal reliable number required for successful queen rearing.

The collective decision of the bees to participate in queen rearing is dynamic, with bees adjusting their behavior to changing conditions. For example, eight bees can successfully rear a queen from a four-day-old larva (34), and even a single worker can care for a four-day-old larva (35). However, fifty bees are insufficient to rear a queen from a one-day-old larva. Interestingly, in half of the cases, the bees began the rearing process and accepted the larva, but later abandoned it. This suggests that the decision to rear a queen depends on a complex interaction between the number of workers and the age of the larva. How the workers assess their number in the cage is a puzzling question that should be further studied. Based on the findings of our first experiment, we continued to investigate the parameters affecting queen rearing in groups of two hundred bees, a practical and easily replicable method. As our protocol became more refined, our success rate in queen-rearing increased in the next experiments.

In our second experiment, we tested the effect of larval age on the quality of the developed queen using two hundred bees in each cage. Several studies have shown that the age of the larvae at grafting affects the quality of the developing queens. Younger larvae tend to develop into higher-quality queens in traditional rearing in foster colonies (15, 19, 33, 36, 37) and in in vitro hand-rearing (25). Our findings support these studies, as we found that larvae aged 0-12 hours developed into heavier queens compared to older larvae. We also demonstrated that the acceptance rate of larvae depends on their age, with larvae older than 48 hours rarely developing into queens (19). The findings of this experiment provide a proof of concept for the new laboratory method of queen rearing, as we were able to consistently replicate former results.

The nutrition available to the workers also influences their decision to accept larvae for queen rearing. When we removed all pollen from the cages, the bees did not accept any of the larvae. A low-protein diet resulted in a low success rate in queen rearing, with only well-nourished workers accepting the task of queen rearing. This finding supports the hypothesis that the nutritional state of the bees is a crucial factor in their decision to rear a queen. The nutritional condition of the workers affects many functions of the bees, including the queen’s egg-laying capacity (22), learning and memory (38) foraging behavior (39) as well as their physiology and immunity (40). Our new method can also be used to test the effects of other nutritional factors on the success of queen rearing, such as different types of pollen, pollen supplements, and pesticide residuals (41-43).

Queen rearing is a major tool for bee stock selection and improvement. The lack of control over queen-rearing conditions has turned this process into a “black box,” where the breeder’s influence on the queen’s development has been limited. Rearing queens under artificially controlled environmental conditions can help extend the queen-rearing season (44). Additionally, isolating developing queens from the hive environment can reduce the impact of pathogens, such as the black queen cell virus, on the success of queen rearing (45). This method can also be employed to test the effects of new pesticides on honey bee health (46, 47). We anticipate that further studies using this new method will shed light on queen development and worker-rearing behavior.

## Acknowledgments

We want to thank the Volcani Institution for providing the facilities and funding for this study. We thank Dr. Victoria Soroker for help with the experiment design and the beekeeper Assaf Otmy for help with the bees.

